# Secondary contact, introgressive hybridization and genome stabilization in sticklebacks

**DOI:** 10.1101/2023.08.29.555285

**Authors:** Xueyun Feng, Juha Merilä, Ari Löytynoja

## Abstract

Advances in genomic studies have revealed that hybridization in nature is pervasive and raised questions about the dynamics of different genetic and evolutionary factors following the initial hybridization event. While recent research has proposed that the genomic outcomes of hybridization might be predictable to some extent, many uncertainties remain. With comprehensive whole-genome sequence data, we investigated the genetic introgression between two divergent lineages of nine-spined sticklebacks (*Pungitius pungitius*) in the Baltic Sea. We found that the intensity and direction of selection on the introgressed variation varied across different genomic elements: while functionally important regions had experienced reduced rates of introgression, promoter regions showed enrichment. Despite the general trend of negative selection, we identified specific genomic regions that were enriched for introgressed variants and within these regions, we detected footprints of selection, indicating adaptive introgression. We found the selection against the functional changes to be strongest in the vicinity of the secondary contact zone and weaken as a function of distance from the initial contact. Altogether, the results suggest that the stabilization of introgressed variation in the genomes is a complex, multi-stage process involving both negative and positive selection. In spite of the predominance of negative selection against introgressed variants, we also found evidence for adaptive introgression variants likely associated with adaptation to Baltic Sea environmental conditions.

## Introduction

Introgression is a process that transfers genetic variation between divergent lineages. Whole-genome analyses have shown introgression to be an important and pervasive evolutionary force (e.g. Mallet, 2005; Lamichhaney et al., 2018; Oziolor et al., 2019) that has shaped the genome of many organisms (e.g. Dowling & Secor, 1997; Harrison & Larson, 2014; Suarez-Gonzalez et al., 2018), including humans (Huerta-Sánchez et al., 2014; Sankararaman et al., 2014, 2016). There is also well-documented evidence of introgression fueling adaptation in several species (Hedrick, 2013; Marques et al., 2019; Racimo et al., 2015; Wang et al., 2023). However, amidst this process of introgression of adaptive and neutral variants, there exists a genome-wide selection against hybrids (e.g. Arnegard et al., 2014; Christie & Strauss, 2018) and regions derived from hybridization throughout the entire genome (Sankararaman et al., 2014; Harrison & Larson, 2014; Juric et al., 2016; Schumer et al., 2018; Calfee et al., 2021). The seemingly conflicting observations of widespread hybridization in nature and the prevalence of selection against foreign ancestry can be explained by the various factors contributing to the reduced fitness of hybrids: In addition to ecological selection against hybrids, the hybridizing parental populations may carry harmful variants (hybridization load), or the genes of the two parental lineages may have negative interactions (hybrid incompatibilities) (Moran et al., 2021).

Genomic studies of contemporary hybrids have shown that the proportion of foreign ancestry is highly variable among species (Martin et al., 2015, 2019; Malinsky et al., 2018) and populations (Skoglund et al., 2015; Kuhlwilm et al., 2016), and the introgressed ancestry is unevenly distributed across the genome (Sankararaman et al., 2014, 2016; Vattathil & Akey, 2015; Zhang et al., 2016). However, the mechanisms underlying this heterogeneous distribution are not well understood. Generally, introgressed alleles are regarded to have a negative fitness effect when introduced into new genomic backgrounds (Amorim et al., 2017; Martin & Jiggins, 2017; Bay et al., 2019) and, according to the Dobzhansky-Muller model of hybrid incompatibility (Bomblies et al., 2007; Masly & Presgraves, 2007; Lee et al., 2008), long-term negative selection on incompatible loci may create “deserts” of introgression in the genome (Sankararaman et al., 2014, 2016). However, genetic architecture and constraints also play a role, and genomic regions characterized by higher gene density and/or low recombination rate are expected to show a lower rate of introgression than other regions (Martin & Jiggins, 2017). This prediction is well supported by empirical studies in humans (Sankararaman et al., 2014, 2016), fish (Schumer et al., 2018), and butterflies (Edelman et al., 2019; Martin et al., 2019), and the intensity of selection against introgression in these studies appears to be positively correlated with the density of functional elements. Nevertheless, the intricate interactions between the different forces against foreign ancestry and the predictability of the ultimate outcomes of hybridization are yet to be fully comprehended.

Introgression and admixture always happen in an evolutionary context, and the hybridizing lineages can differ in terms of their demographic histories and population sizes, levels of genetic drift, and strength of selection against deleterious variants prior to the hybridization event (Schumer et al., 2018; Moran et al., 2021; Liu et al., 2022). Gene flow from a population with a smaller effective population size (*N_e_*) and reduced purifying selection efficiency may increase the genetic load in the hybrid population through the introduction of weakly deleterious alleles (Harris & Nielsen, 2016; Juric et al., 2016). Typically, gene flow from a population with a larger *N_e_* is thought to ease the genetic load (Edelman & Mallet, 2021) but, in extreme cases, it may import unbearable amounts of recessive lethal variation and condemn a tiny population into extinction (Kyriazis et al., 2021). Consequently, the selection on introgressed variants, both adaptive and maladaptive, plays a pivotal role in shaping the genome-wide patterns of foreign ancestry (Kim et al., 2018). The simultaneous operation of multiple demographic and selective processes may lead to interference effects, emphasizing the need for a systematic investigation of the historical demographic events and the distinct evolutionary forces that shape the genomic landscape of introgression. A thorough understanding of both the history and the genomic mechanisms is vital for comprehending the evolutionary consequences of hybridization.

The nine-spined stickleback (*Pungtitius pungitius*) is a small euryhaline teleost fish that inhabits circumpolar regions of the northern hemisphere. The evolutionary history of nine-spined sticklebacks has been extensively studied (Aldenhoven et al., 2010; Shikano, Ramadevi, et al., 2010; Teacher et al., 2011; Bruneaux et al., 2013; Natri et al., 2019; Guo et al., 2019; Feng et al., 2022, 2023), and in Europe two distinct evolutionary lineages have been identified: the Western European lineage (WL) and the Eastern European lineage (EL). Despite the lineages exhibiting distinct sex determination systems (Natri et al., 2019) and highly differentiated mitochondrial haplotypes (Aldenhoven et al., 2010; Shikano, Shimada, et al., 2010; Guo et al., 2019), they are known to interbreed (Natri et al., 2019) and populations in the Baltic Sea area display a gradient of mixed ancestry (Feng et al., 2022). The identity of the participating populations and the exact timing of the Baltic Sea admixture event(s) are unknown. Based on purely geographical information (Ukkonen et al., 2014), the EL appears to have colonized the Baltic Sea relatively late (in the Ancylus Lake stage, 10,700–10,200 BP; Björck, 2008; Feng et al., 2022) and it seems likely that the large water body had (WL-origin) sticklebacks trapped for the duration of the ice age or at least the species had colonized the area during the Yoldia Sea stage (11,600–10,700 BP). If the area was inhabited by WL-origin sticklebacks, the two lineages met the first time when the EL colonized the Baltic Sea; the second window for admixture started after the re-opening of the Danish Straits during the Littorina Sea stage (at the latest 8500–6000 BP). If the pre-Ancylus waters were void of sticklebacks, the area was first of EL-origin and the two lineages have admixed during or after the Littorina Sea stage and the formation of the current Baltic Sea.

The Baltic Sea is a relatively shallow brackish-water inland sea with steep gradients of salinity and other abiotic factors (Szaniawska, 2018; Garcia et al., 2019). Due to its young age and recent colonization by various species, it provides an appealing system to study adaptation to variable conditions (Reusch et al., 2018). The genetic resources, the broad-scale sampling, and the detailed demographic history available for the Baltic Sea nine-spined sticklebacks (Varadharajan et al., 2019; Kivikoski et al., 2021; Feng et al., 2022) provide a good starting point for separating the complex signals of various evolutionary factors. The known genetic and phenotypic variation in the area (Herczeg et al., 2009, 2010; Shikano, Ramadevi, et al., 2010; Teacher et al., 2011; Guo et al., 2019; Natri et al., 2019; Feng et al., 2022, 2023) make the species an intriguing model for investigating admixture, and the presence of populations with large *N_e_* reduces the impact of drift (Palumbi, 1994; Tigano & Friesen, 2016; Jokinen et al., 2019) and the hybridization load and thus permits excluding specific mechanisms from the process. Ideally, the Baltic Sea sticklebacks should allow studying the consequences of admixture and the stabilization of the post-admixture genome and help understand the interplay of different forces and mechanisms that lead genomes to resist ongoing introgression in hybrid zones.

The objective of this study was to investigate the mechanisms influencing the introgression landscape between two evolutionary lineages of nine-spined sticklebacks (*Pungitius pungitius*) after their secondary contact in northern Europe. To this end, we employed whole-genome sequence data from 284 individuals belonging to 13 populations across the Baltic Sea hybrid zone, subsampled from Feng et al. (2023). Our analysis aimed to estimate the levels of introgression across various populations and different genomic regions, thereby characterizing the genomic landscape of introgression across the hybrid zone. Our main objectives were threefold: 1) to comprehend the leading evolutionary forces that have shaped the genomic landscape of introgression, 2) to identify genomic regions where positive natural selection has likely favored introgressed genetic elements, and 3) to elucidate how natural selection has influenced the genome-wide patterns of introgression, particularly regarding the transfer and elimination of adaptive and deleterious variants between the two lineages. Our findings suggest that a diverse array of evolutionary forces has contributed to shaping the genomic landscape of introgression, the fast removal of highly deleterious variations and the long-term selection against weak deleterious variations being the predominant driving forces. The different sources of selection have interacted with the variable recombination landscape and genome structure, thereby adding complexity to the predictability of post-admixture genome evolution in the hybrids.

## Materials and Methods

### Ethics Statement

The data used in this study were collected in previous studies and in accordance with the national legislation of the countries concerned.

### Data Acquisition

The data were subsetted from the vcf file provided in Feng et al. (2023) using BCFtools v.1.7 (Li et al., 2009). Following Feng et al. (2022), DEN-NOR (from the North Sea) was used as the WL source population, GBR-GRO (from the UK) as the WL reference population, RUS-LEV (from the White Sea) as the EL source population, and CAN-TEM (from Quebec, Canada) was selected as the outgroup. As the focal study populations, nine admixed marine populations from the Baltic Sea identified by Feng et al. (2022) were selected (Fig. 1a and Table S1). In the earlier analyses of this data (Feng et al., 2023), the reads were first mapped to the latest nine-spined stickleback reference genome (Kivikoski et al., 2021) using the Burrows-Wheeler Aligner v.0.7.17 (BWA-MEM algorithm, Li, 2013) and its default parameters. Duplicate reads were marked with SAMtools v.1.7 (Li et al., 2009) and variant calling was performed with the Genome Analysis Toolkit (GATK) v.4.0.1.2 (McKenna et al., 2010) following the GATK Best Practices workflows. Sites located within identified repetitive sequences (Varadharajan et al., 2019) and negative mappability mask regions combining the identified repeats and unmapped regions (Kivikoski et al., 2021) were excluded. Multiallelic variants and sites showing an extremely low (< 5x) or high average coverage (> 25x), genotype quality score (GQ) < 20, quality score (QUAL) < 30 and missing data > 75% were filtered out using VCFtools v.0.1.5 (Danecek et al., 2011). Data from the known sex chromosomes (LG12, Natri et al., 2019) were removed from further analysis.

**FIGURE 1.**
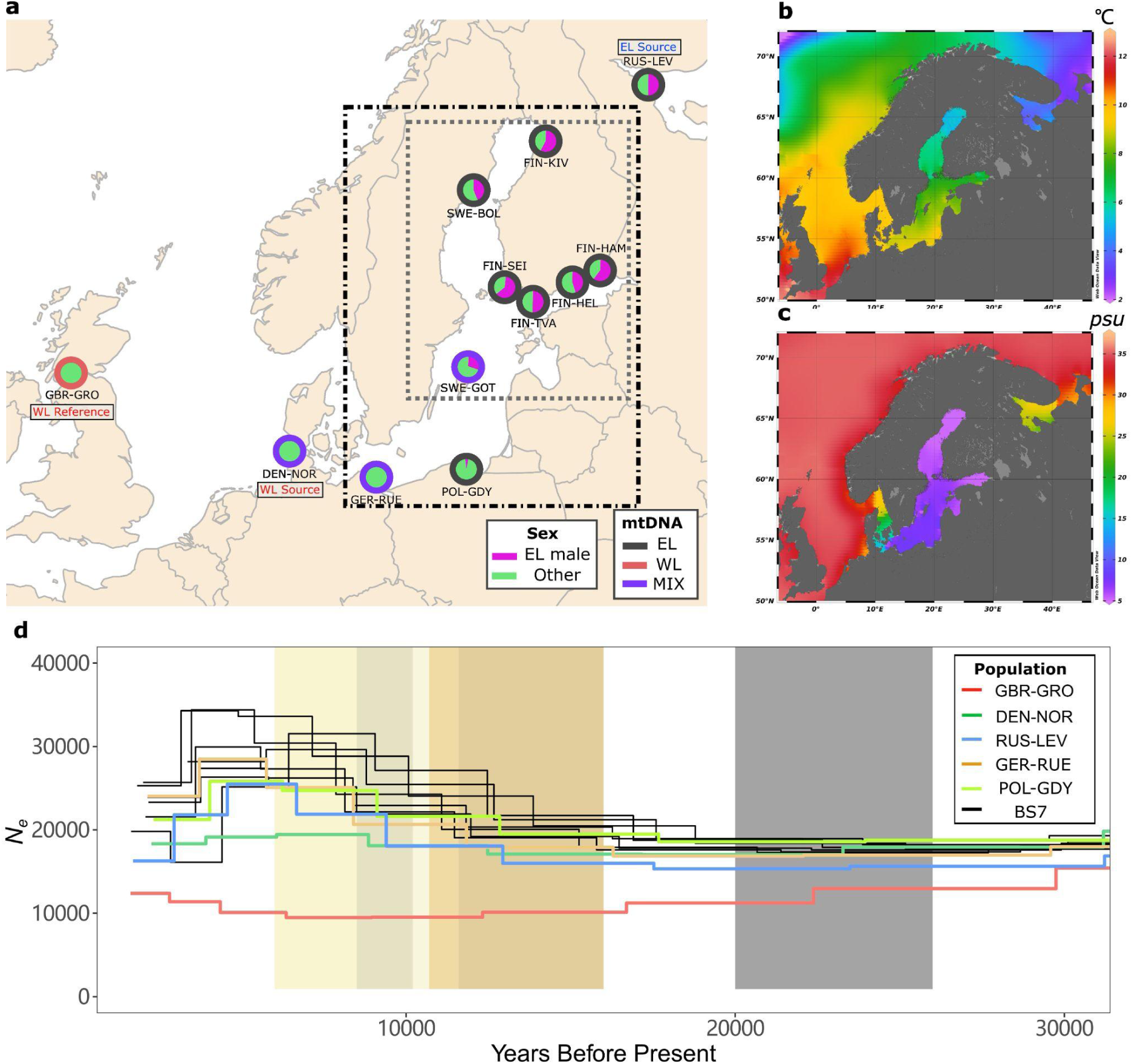
Study populations and localities. (a) Geographic origins of populations involved in this study, modified from Feng et al. (2022). The pie charts show the proportions of Eastern Lineage (EL) males; “Other” can be either Western Lineage (WL) male or WL/EL female. The outline colors (black, red, and purple) indicate the mtDNA lineage assignment of the population. The dot-dashed frame marks the admixed populations and the gray dotted frame indicates the BS7 set (see Methods). The source and reference populations used in the *f4-ratio* test and *fd* statistics are indicated. (b) Map of the sea surface temperature and (c) salinity of the Baltic Sea and its surroundings, adapted from Garcia et al. (2019). (d) Demographic history of parental, reference, and admixed populations from 30,000 years ago (kya) to the present, adapted from Feng et al. (2023). The blue, green, and red colors indicate the EL parental population RUS-LEV, WL parental population DEN-NOR, and WL reference population GBR-GRO, respectively; the orange and green colors indicate GER-RUE and POL-GDY, the two southern Baltic Sea populations, respectively. The seven populations from the central and northern Baltic Sea are depicted in black. The yellow shadings indicate different stages of the Baltic Sea: the Baltic Ice Lake (16,000–11,600 BC), the Yoldia Sea (11,600–10,700 BC), the Ancylus Lake (10,700–10,200 BC), the fresh-to-brackish water transition stage (10,200-8,500), the Littorina Sea (8,500–6,000 BC). The gray shading indicates the last glacial maximum (26,000–20,000 BC).

### Quantification of Genomic Introgression

The f_4_-ratio test (Reich et al., 2009) was applied to quantify the amount of foreign ancestry in different genomic features. Following Petr et al. (2019) and Feng et al. (2022), the WL ancestry (*α*_WL_) was estimated as:

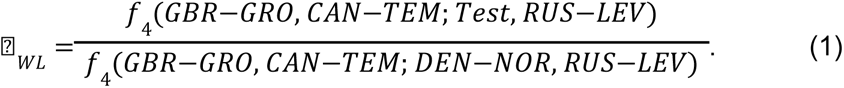

*f_4_-ratio* tests were performed using ADMIXTOOLS v.5.1 (qpDstat v.755, qpF4ratio v.320; Patterson et al., 2012).

To estimate the admixture proportion among different genomic features, the locations of constrained elements were lifted from the three-spined stickleback genome annotation (Ensembl release ver. 95; Herrero et al., 2016) using CrossMap v.0.3.3 (Zhao *et al*. 2014) and a liftover-chain created with LAST (Frith et al., 2010) and the Kent utilities (Tyner et al., 2017). The genome annotations from Varadharajan et al. (2019) were lifted to the latest nine-spined stickleback reference genome (Kivikoski et al., 2021) using liftoff (Shumate & Salzberg, 2021). The promoter regions were defined as 1kb stretches upstream of the gene start. A significance test was then applied to assess whether EL and WL ancestry were significantly enriched or depleted in any of the genomic features in comparison to the levels seen within intergenic regions. Following Petr et al. (2019), the alpha value of a given annotation category was resampled 10,000 times from a normal distribution centered on the alpha with a standard deviation equal to the standard error given by ADMIXTOOLS. An empirical *p*-value was then calculated for the estimated alpha for each genomic feature to test the hypothesis that the ancestry proportions for different genome features do not differ from that of the intergenic regions.

### Quantification of WL Introgression (*fd*) and Population Genetic Statistics

The modified D-statistic, *fd* (Martin et al., 2015), was used to quantify introgression for the admixed population at finer genomic scales. We used a fixed window size of 100 kb with a 20-kb step size using the scripts from Martin et al. (2015) and estimated *p*-values from the *Z*-transformed *fd* values using the standard normal distribution and corrected for multiple testing with the Benjamini-Hochberg false discovery rate (FDR) method (Benjamini & Hochberg, 1995). Windows with positive *D* and *fd* values with a number of informative sites ≥ 100 and FDR value ≤ 0.05 were retained as outlier loci (see below). Similarly to the *f4*-ratio test, we used the DEN-NOR and RUS-LEV to represent the WL and EL ancestral population and CAN-TEM as the outgroup. The admixture proportions were estimated separately for each Baltic Sea population and for a combined set of seven northern populations showing similar levels of introgression (i.e., SWE-GOT, FIN-HEL, FIN-TVA, FIN-HAM, FIN-SEI, SWE-BOL and FIN-KIV, hereafter BS7 when referred collectively).

We examined the covariation of admixture proportions (*fd*) with the population genetic statistics *π* (nucleotide diversity), *d_xy_*(absolute divergence), and *F_ST_* (measure of genetic drift). The statistics were computed genome-wide in 10 and 100-kb windows using the scripts from Martin et al. (2015). The mean recombination rate was estimated from the linkage map Varadharajan et al., (2019), initially for 10-kb windows (see Varadharajan et al., 2019 for details) and then binning the rates into 100-kb non-overlapping windows. The rates were lifted to the latest nine-spined stickleback reference genome (Kivikoski et al., 2021) using custom scripts. The 10-kb population genetic statistics were used in fine-scale analyses of candidate regions for adaptive evolution.

### Footprints of Selection in Baltic Sea Populations

After the quantification of WL introgression, we searched for footprints of selection on introgressed variants using the U and Q95 tests following Racimo et al. (2016) and Jagoda et al. (2018). Both tests are based on the variant allele frequencies (VAF) and measure, respectively, the number of single-nucleotide polymorphisms (SNPs) shared with the donor population that appear at a high frequency in the focal population but at a low frequency in the reference population, and the 95% quantile of the frequency of the SNPs that are shared with the donor population and appear at a low frequency in the reference population. The VAF was estimated separately for WL (DEN-NOR), EL (RUS-LEV), and the Baltic Sea populations, and the tests were performed using 100-kb windows with a 20-kb step and discarding positions with more than 25% missing data. We first calculated the *U20*_EL, BALTIC, WL_(0.01, 0.2, 1) to count the SNPs that are at < 1% frequency in the combined EL reference population, at ≥ 20% frequency in the combined Baltic Sea population (BS7), and fixed (100% frequency) in the WL population. We then calculated the *Q95*_EL, BALTIC, WL_(0.01, 0.2, 1) to obtain the 95% quantile VAF of these SNPs in the Baltic Sea population. The intersection of the top 1% regions of the *U20* and *Q95* tests and the candidate regions from the *fd* test were then considered as putative adaptive introgression (AI) regions. Within each AI region, *F_ST_*, *d_xy_* and *π* were calculated for 10-kb sized windows with a 5-kb step, and genotypes and allele frequencies (minor allele frequency [*maf*] ≥ 0.05) of variants were used to identify candidates for possible adaptive evolution among the lifted reference gene annotations.

### Assessment of Introgressive Genetic Load and Its Purification in the Baltic Sea

Following the concept of U20 test, we defined the variants in the Baltic Sea populations to be of WL-origin if they showed frequency ≤ 0.05 in the EL reference and frequency ≥ 0.95 in the WL reference. To evaluate the efficacy of selection on potentially deleterious foreign variations introduced via genetic introgression, we employed the r_xy_ statistics as described by Xue et al. (2015). The r_xy_ statistics compared the allele frequencies of certain categories of variants relative to the levels of neutral variants between populations located at varying distances from the entry to the Baltic Sea. An r_xy_ value below 1 indicates a deficiency of the focal alleles in the population farther from the WL introgression entry, suggesting the influence of purifying selection. The effects of variants on protein-coding gene sequences were annotated and classified as low impact (synonymous variants), moderate impact (missense variants), and high impact (stop codon gaining variants) using SnpEff v.5.0 (Cingolani et al., 2012). The same number of variants from intergenic regions were randomly selected and served as a proxy for the neutral level of genetic variation. We obtained standard errors and 95% confidence intervals for the r_xy_ estimates by jackknifing the values across the 20 individual linkage groups (LG).

## Results

### Data Description

We retained 284 individuals from 13 populations of the total data of 888 individuals and 45 populations studied earlier by Feng et al. (2022 & 2023) (Fig. 1a and Table S1). The admixed populations were required to be from marine localities with variation in environmental conditions (Fig. 1b and c) and have large *N_e_* (Fig. 1d; Feng et al., 2023) to minimize the impact of genetic drift. The reference populations for the *f4-ratio* test and *fd* statistics were selected based on Feng et al. (2022). After quality control, 4,294,816 biallelic SNPs over the 20 autosomal LGs remained for the following analyses.

### Quantification of Introgression Across the Genome

Following Petr et al. (2019) we applied the *f_4_-ratio* test (Reich et al., 2009) to quantify the composition of WL ancestries in the Baltic Sea populations. In the intergenic regions, considered representing the background level, the southern Baltic Sea populations (GER-RUE and POL-GDY) contain 35% and 22.23% of WL ancestry (Fig. 2), respectively, whereas the more northern populations from Gotland (SWE-GOT), Gulf of Finland (FIN-HEL and others) and the Bothnian Bay (SWE-BOL, FIN-KIV) contain 13.48–11.27% of WL ancestry, consistent with the whole-genome estimates of Feng et al. (2022). Overall, the ancestry proportions show opposite gradients across the Baltic and North Seas, with the WL ancestry decreasing with increasing distance from the Danish straits (Fig. 2, Supplementary Table S2).

**FIGURE 2.**
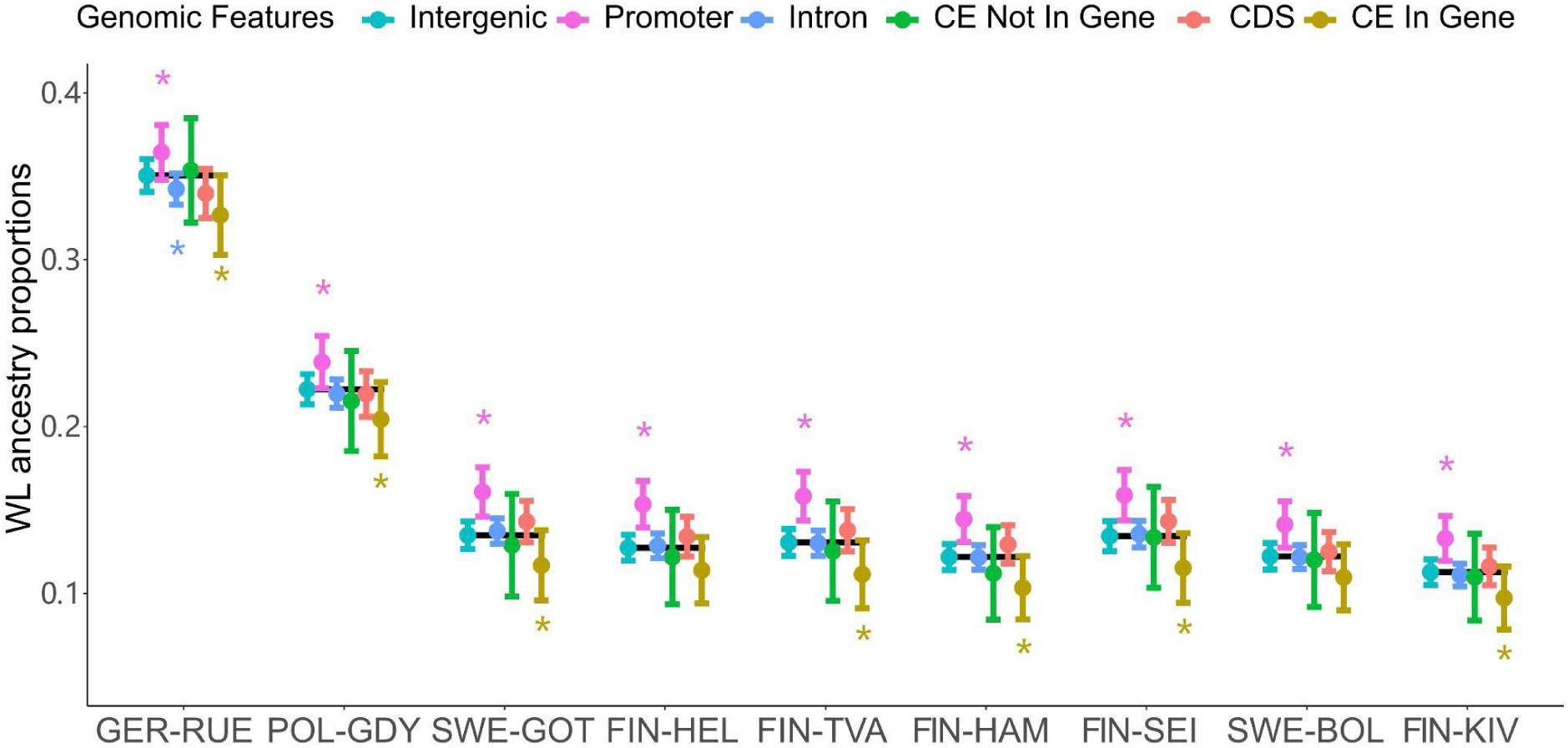
Western Lineage ancestry across six different genomic features in the admixed Baltic Sea populations. The dots and lines show the proportion of WL ancestry and its standard error (estimated with jack-knifing across the genome) for each genomic feature, the asterisks above and below indicating that the estimate is significantly higher or lower than within the intergenic region. The black lines indicate the estimates for intergenic regions, assumed to represent the background level of WL ancestry. The populations are ordered by increasing geographic distance from the Danish Straits. CE = constrained elements, CDS = coding sequences.

To examine the potential impacts of selection on the minor parental ancestries, we binned the genome into functional categories and computed the WL ancestries across them using the *f_4_-ratio* test. We considered six categories: intergenic, coding DNA (CDS), constrained elements located inside or outside of genes, introns, and promoters. According to these estimates, all Baltic Sea populations contain significantly elevated amounts of WL ancestry in promoter regions (*p* < 0.001–0.044, estimated via resampling, see Methods; Supplementary Table S2). Additionally, except for FIN-HEL and SWE-BOL (p = 0.090–0.098), all populations display significantly lower amounts of WL ancestry in constrained elements located within genes (p = 0.024–0.055). While not statistically significant for all mid to northern Baltic Sea populations, there was a slight increase in WL ancestry in CDS compared to intergenic regions.

### Footprints of Selection on WL Introgression in Baltic Sea Populations

The seven populations from the northern Baltic Sea showed similar levels of genetic introgression and were studied more closely to understand the factors shaping the genomic landscape of introgression as well as the potential adaptive nature of the introgressed variation. To more precisely identify the regions enriched with introgressed WL variants, we combined the populations, referred to as BS7, and computed the *fd* summary statistic (Martin et al., 2015) for 100-kb windows with 20-kb steps across the genome. Based on the FDR corrected *p*-value cut-off at 0.05, we obtained 181 putative introgression-enriched regions; by merging the overlapping regions, these collapsed into 45 regions with lengths varying from 100–560 kb (Fig. 3a, Supplementary Table S3).

**FIGURE 3.**
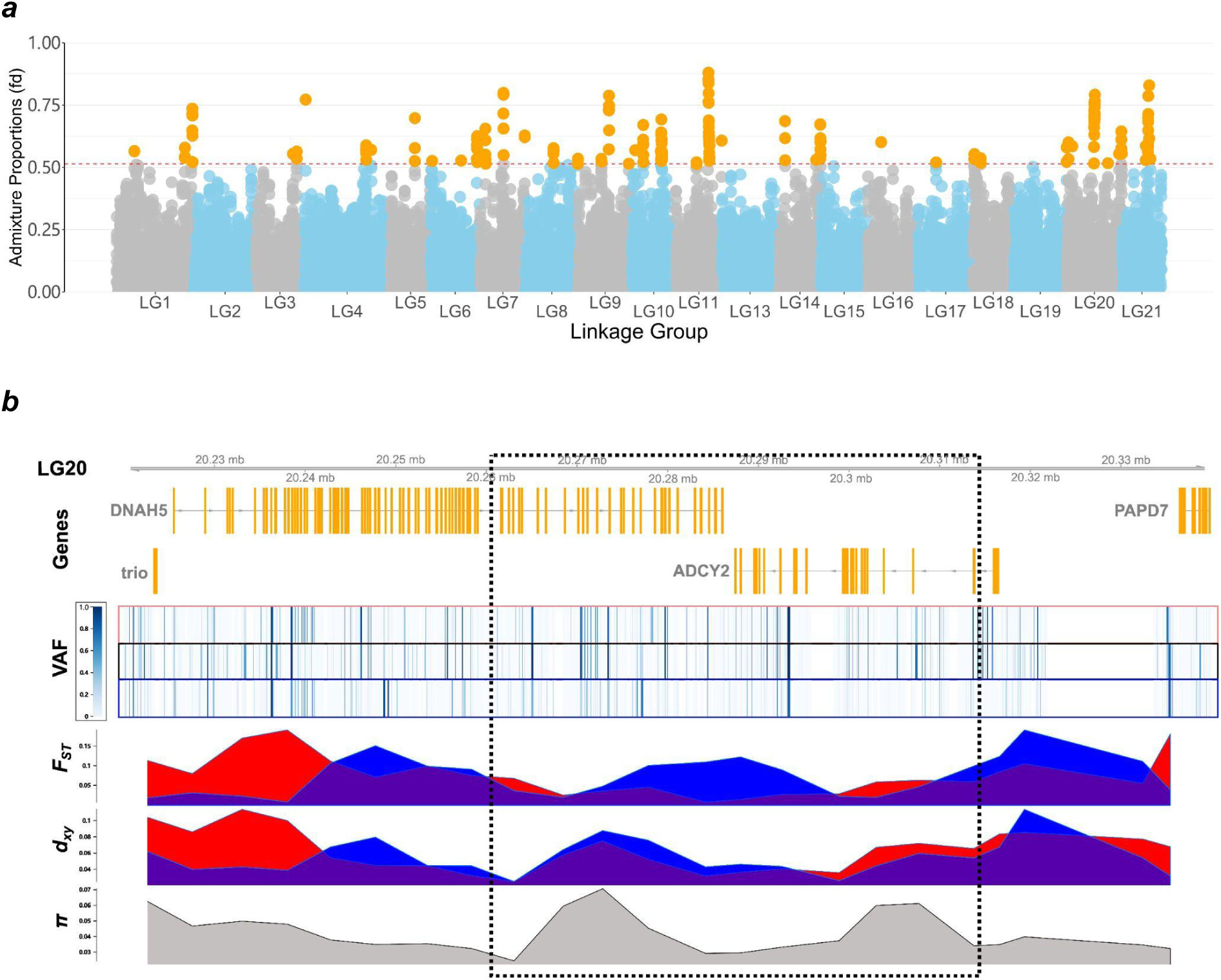
Adaptive introgression in the northern Baltic Sea populations. (a) The Manhattan plot shows the estimated admixture proportions (fd) for 100kb windows across the genome, orange dots indicating genomic windows significantly enriched for WL ancestry. The dotted red line at 0.5137 indicates the *p*<0.05 significance level. (b). A candidate region for adaptive introgression (AI) at LG20:20260000-20315000 (dotted box). The panels show the gene annotations (coding sequences in orange), per site variant allele frequencies (VAF; heatmap) for the three sets of population (WL source, BS7, EL source), and *F_ST_*, *d_xy_* and *π* (10kb-windows) in the northern Baltic Sea populations (BS7). The red and blue colors indicate *F_ST_* and *d_xy_* measured against DEN-NOR (WL source) and RUS-LEV (EL source), respectively.

By definition, introgression introduces novel variation to a population and the footprints of selection within introgressed genomic regions differ from those expected under models without introgression (Setter et al., 2020). More precisely, methods based on polymorphism patterns may fail to detect the signals of selection, and approaches based on the overrepresentation of introgressed alleles in a specific population relative to other populations are considered more robust (Racimo et al., 2015). Following this, adaptive introgression can be distinguished from neutral admixture using the number of sites uniquely shared between the donor and recipient population (*U* test) as well as the allele frequencies on those sites (*Q95* test; Racimo et al., 2016). We applied the *U* and *Q95* tests to search for footprints of selection amongst the introgressed variants and obtained 44 regions which collapsed into eleven candidate regions (Supplementary Table S4). Five of these candidate regions overlapped with the regions identified with the *fd* analysis and were chosen as candidates for adaptive introgression (AI). Integrating information from *F_ST_*, *d_xy_*, variant allele frequencies, and genetic diversity, we identified four candidates for adaptively introgressed genes (*viz*. *ZP4*, *PLEKHG3, DNAH5, ADCY2*; Supplementary Table S4, Fig. S1). The region in LG20 shows all the hallmarks of a selective sweep (Fig. 3b): lowered *F_ST_* to the WL reference and increased *F_ST_* to the EL reference, no significant increase in *d_xy_*, and a volcano-shaped pattern of *π* created by recombination between the alternative haplotypes (Setter et al., 2020). Although the number of genes found within the candidate regions is small, this does not exclude the possibility that a greater number of genes would be under adaptive selection, e.g., through introgressed promoter regions or other regulatory elements and structural variations.

### Interaction between Introgression, Differentiation, and Recombination Rate

Possible mechanisms causing heterogeneity in admixture proportions and differentiation across the recipient populations’ genomes include incompatibility of genetic variants introgressed from a diverged lineage (Schumer *et al*. 2018), selection against introduced deleterious variation (Juric et al., 2016; Kovach et al., 2016) and adaptive evolution in different environments (Chunco, 2014; Smukowski Heil et al., 2019). As selection removes negative variants, it also removes linked neutral variation, giving the variation in recombination rate a role in shaping the distribution and patterns of introgression across the genome (e.g., Schumer et al., 2018). In our data, WL ancestry proportion and recombination were only weakly positively correlated in the German coastal population (*r_s_* = 0.064, *p* < 0.001). Interestingly, no correlation was found in the Polish population (*r_s_* = −0.029, *p* = 0.082) and in Gotland (*r_s_* = −0.023, *p* = 0.174) but a weakly negative correlation was seen in the other mid and northern Baltic Sea populations (*r_s_* = −0.090–−0.039, *p ≤* 0.018). In the combined BS7 set, the correlation was slightly negative (*r_s_* = −0.046, p = 0.006; Table 1).

**TABLE 1.**
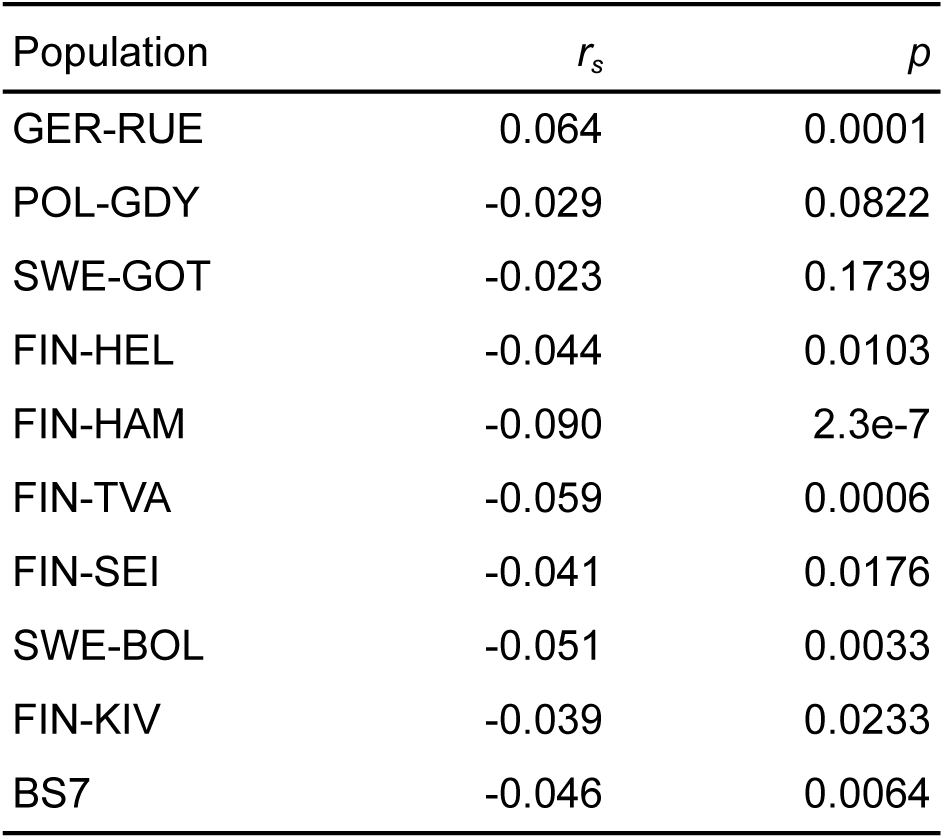
Spearman rank correlations between admixture proportion (*fd*) and recombination rate in different populations. BS7 = combined seven northern Baltic Sea populations.

Due to decreased *N_e_* (and increased drift) caused by background selection, genomic regions experiencing lower levels of recombination are expected to be more differentiated than those experiencing more recombination (Nachman & Payseur, 2012). In our data, the genetic diversity, as well as the genetic distance measurements *F_ST_* and *d_xy_*, regardless of which parental population they were compared with, were always positively correlated with recombination rate (*r_s_*= 0.179–0.615, *p ≤* 0.001; Table S5). As the density of coding sequences across the genome is weakly and positively correlated with the recombination rate (*r_s_* = 0.098, *p* < 0.001), it is not surprising to see the admixture proportion also weakly correlating with the density of coding sequences (*r_s_*= −0.047, *p* = 0.006) and density of constrained elements (*r_s_*= −0.074, *p* < 0.001).

### Introgressive Genetic Load Across Populations

Given the dramatic differences in WL ancestry across the localities and the different genomic features, we set out to investigate the steps, progression, and stabilization of the WL ancestry within the genome since the secondary contact. Specifically, we investigated how the efficacy of selection on introgressed variants changes across the Baltic Sea (Fig. 4a). First, we identified WL-origin variants with opposing allele frequencies (Fig. 4b); these alleles were found in high frequencies in the south but their frequency dramatically decreased in the central Baltic Sea and further declined towards the north. Second, we estimated the *r_xy_* statistics for all coding variants and the WL-origin coding variants. The results show that, while no apparent differences are seen at the genome-wide level (Fig. 4c), the selection has very efficiently purged the introgressed genetic variation in the south compared to the mid and northern Baltic Sea (Fig. 4d), especially the variants inferred to have a significant effect and causing early stop codons (High Impact). Interestingly, we found that the strength of purging of WL-origin variation is not much different between synonymous (Low Impact) and nonsynonymous (Moderate Impact) variants. The efficiency of the purging of WL-origin variation does not appear to depend on the recombination rate (*r_s_*=−0.176, *p*=0.220; Supplementary Fig. S2).

**FIGURE 4.**
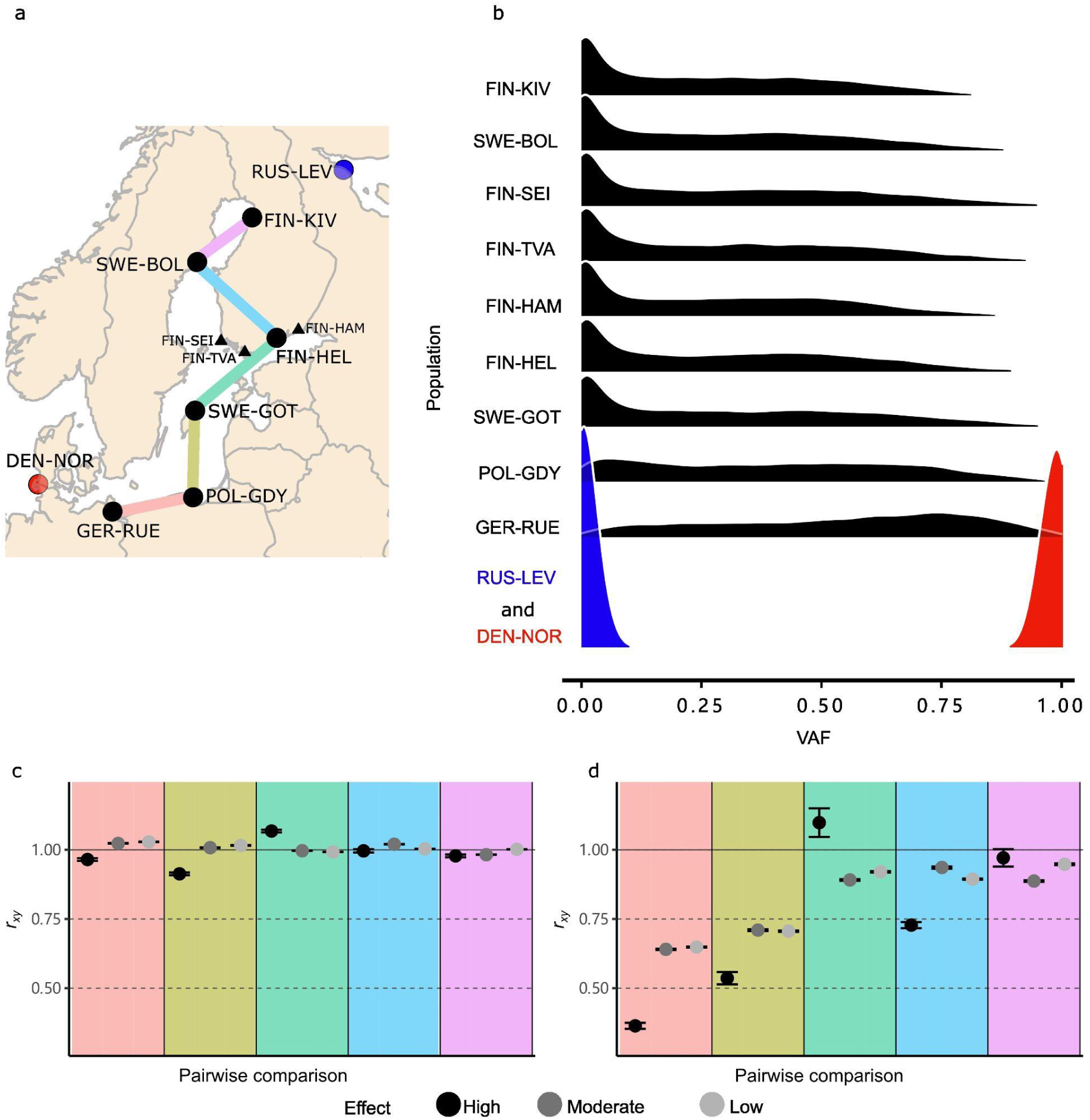
Selection on potentially deleterious introgressed variation. (a) Populations for the pairwise *r_xy_* comparisons with colors matching those in panels c and d. The three populations from the Gulf of Finland indicated with black triangles were not included in the pairwise comparisons. The populations representing the WL and EL parental lineages are indicated with red and blue dots, respectively. (b) Density plots of the WL-origin variant allele frequencies (VAF) in the admixed Baltic Sea populations and the two parental populations. (c) and (d) The *r_xy_* statistics for all (c) and WL-origin (d) coding variants of different impact for the five pairwise comparisons (Supplementary Table S6). *r_xy_* =1 if the allele frequency changes do not deviate from the background; *r_xy_* < 1 indicates the relative deficit of the corresponding variants in the northern population compared to the southern population. *r_xy_* confidence intervals are based on jack-knife estimates across the 20 linkage groups.

## Discussion

In hybridization, two locally adapted genomes get mixed and the intra-genomic interactions get broken, opening a window for exceptionally dynamic evolution followed by a phase of subsequent stabilization. Large-scale DNA sequencing has revealed the prevalence of genetic introgression in the wild, but the events after the hybridization and the roles and the interplay of the different evolutionary factors are more difficult to study and still poorly understood. While some basic principles of hybridization have been emerging (Moran et al., 2021) and it is well documented that the foreign ancestry is selected against within the functionally most important genome regions, little is known about the relative importance of the different evolutionary forces and their interactions, and ultimately, how predictable the outcomes of hybridization are.

We investigated the genome-wide patterns of genetic introgression between two divergent lineages of nine-spined sticklebacks across the Baltic Sea and found the minor parental ancestry being generally selected against, with only a few regions showing signals of selection favoring the foreign ancestry. We found little correlation between the admixture proportion and the recombination rate, indicating a limited role for recombination in shaping the genomic landscape of introgression in this model system. Although we cannot fully separate the signals created by the temporal factors of a potentially multi-staged admixture history from those created by the selection, the results highlight the complex forces acting in hybrid populations and demonstrate the potential of the Baltic Sea sticklebacks as a model system to study the evolutionary processes after secondary contacts in action.

### Genomic variation of minor parental ancestry

In hybrid populations, genome-wide patterns of ancestry are predicted to be extremely variable right after the hybridization event and then gradually stabilize over time. In our study, the WL ancestry decreased from 35% in the population closest to the entry to the Atlantic to 22.2% in a population 330 km to the east and then leveled at 13.5–11.3% in mid and northern Baltic Sea populations. In principle, this could be explained by the ∼12% WL ancestry in the mid and northern parts of the Baltic Sea representing the ancestral, first-stage admixture event, and the higher WL ancestry levels in the south being explained by more recent and still ongoing WL migration. However, consistent with a number of previous studies showing that hybridization is selected against (Arnegard et al., 2014; Sankararaman et al., 2014; Juric et al., 2016; Harris & Nielsen, 2016; Christie & Strauss, 2018; Schumer et al., 2018; Calfee et al., 2021), we found that the selection has specifically removed the functionally significant WL ancestry in the southern Baltic Sea, both in the most conserved parts of coding sequences (Fig. 2) and among functional coding sequence changes (Fig. 4d). While the genome-wide ancestry proportions in the mid and northern parts of the Baltic Sea are highly similar, there is a consistent pattern of WL ancestry being enriched in the promoter regions and being depleted in the constrained elements within coding sequences. Consistent with the latter, the *r_xy_* values for the amino-acid-changing variants of WL-origin are below one in all pairwise comparisons, indicating selection against the functionally significant foreign variation. Similar patterns of rapid removal of largely deleterious introgressed variation have been found in studies of hominins, swordtail fishes and *Drosophila* (Harris & Nielsen, 2016; Schumer et al., 2018; Matute et al., 2020; Veller et al., 2023), and such a pattern is considered to be widespread.

### Diverse Evolutionary Forces Shaped the Landscape of Introgression

Recombination rate variation is known to play a key role in shaping the genomic landscape of introgression (e.g. Kim et al., 2018; Martin et al., 2019; Veller et al., 2023). Generally, the footprints of selection are more prominent in regions with low recombination rates where the minor parental haplotypes are longer and thus more likely to contain harmful alleles (Wu, 2001; Nachman & Payseur, 2012). As a result, a positive correlation between the admixture proportion and recombination rate is expected if the selection against genomic incompatibilities is the dominant force (e.g. Schumer et al., 2018; Martin et al., 2019; Edelman et al., 2019); conversely, a negative correlation is expected if the foreign ancestry is favored (Pool, 2015; Corbett-Detig & Nielsen, 2017; Duranton & Pool, 2022) or if the effects of deleterious variants are recessive (Kim et al., 2018). Interestingly, we found that the correlation between admixture proportions and recombination rates was in general weak and varied across the Baltic Sea: correlation was slightly positive in Germany, non-existent in the central Baltic Sea, and weakly negative in the more northern parts. The pattern is consistent with a model where the selection against introgression varies during the process. At the early stages, the selection against incompatibilities and highly deleterious variation is the dominant force, creating a positive correlation; with increasing distance from the secondary contact zone, the foreign ancestry gets “filtered” and the force of selection against it diminishes, which in turn weakens the correlation between the recombination rate and the levels of introgression (Groh & Coop, 2023).

Consistent with this, we found that the purifying selection has very efficiently purged early stop codons and amino-acid changing variation of WL-origin in the southern and central parts of the Baltic Sea, while the selection against the different types of variants tends to become more similar towards the north (Fig. 4). Importantly, at the genome-wide level, we found no evidence of purging of the putatively deleterious moderate and low impact variants, and thus neither drift nor selection has significantly affected the overall proportions of weakly deleterious alleles. The deficit of high-impact coding variants of WL origin in the southern Baltic Sea is surprisingly strong (Fig. 4); as the WL-origin variants appear at a frequency >0.95 in the North Sea population, they cannot be highly deleterious in that environment. The strong selection against them indicates that they must be maladaptive on the Baltic Sea side and suggests that the environmental differences are the driver of this selection.

The observed enrichment of introgression within promoter regions (Fig. 2) is contrary to findings in human studies (Petr et al., 2019; Telis et al., 2020) but agrees with those in swordtail fishes (Schumer et al., 2018). In swordtail fishes, recombination is concentrated within promoter and other functional regions (Baker et al., 2017), whereas in humans the process is driven by specific DNA motifs detected by the chromatin-modifying protein PRDM9 (Myers et al., 2010). The relationship between recombination and genomic features in the nine-spined stickleback is unknown, and recombination cannot be excluded as a factor explaining the high WL ancestry within promoters. On the other hand, introgression is more likely to induce regulatory changes in gene expression than radical alterations in protein-coding genes (Gittelman et al., 2016; Dannemann et al., 2017; Dannemann & Kelso, 2017; McCoy et al., 2017; but see Petr et al., 2019; Telis et al., 2020). If so, the strong and consistent pattern across all populations could suggest that a selective sweep happened in the first admixture event between the EL and the ancestral Baltic Sea population. The opposite (and consistent) trends for the classes “Promoter” and “CE In Gene” in WL ancestry are striking and may suggest that genetic incompatibilities take place on the level of coding genes (selecting against minor parental ancestry; here WL) while the adaption is driven by gene regulation (favoring the local ancestry; here ancestral Baltic Sea of WL origin).

### Adaptive Introgression

Recent studies suggest that introgression is an important source of genetic variation and allows adaptive evolution to proceed much faster than it would do with *de novo* mutations (Racimo et al., 2015; Edelman & Mallet, 2021). Evidence for adaptive introgression has been found in a diverse array of taxa, including humans (e.g. Racimo et al., 2015), fish (e.g. Meier et al., 2017; Oziolor et al., 2019), butterflies (The Heliconius Genome Consortium, 2012; Pardo-Diaz et al., 2012), and plants (Whitney et al., 2006). Our results add to this evidence by identifying four genes with high-frequency WL-origin variants in the EL genetic backgrounds with footprints of selection.

Two of these genes might be associated with reproduction. The zona pellucida glycoprotein 4 (ZP4) gene is known for its importance in sperm-egg interactions during fertilization (Wassarman et al., 2001). As a key component of the fish chorion (egg cell coat), it could have a role in the adaptation to the brackish water conditions of the Baltic Sea (Lønning & Solemdal, 1972; Nissling, 2002; Jovine et al., 2005). The second gene, dynein axonemal heavy chain 5 (DNAH5), encodes dyneins, essential for flagellar beating and sperm function (Turner, 2003). Harmful mutations in the DNAH5 gene have been associated with dysfunction in spermatozoa (Zuccarello et al., 2008). Notably, DNAH5 has also emerged as a candidate gene behind salinity-associated ecological speciation in the Baltic Sea flounder: The footprints of selection in DNAH5 suggest that the gene has contributed to the flounders’ adaptation to low-salinity Baltic Sea conditions and allow sperm activation in lower salinities than in the related saltwater-adapted European flounder (Momigliano et al., 2017; Jokinen et al., 2019).

The relatively small number of positively selected candidates from our study may be an underestimate due to challenges in identifying positively selected introgressed variants and our rather stringent approach to screening for candidates of adaptive introgression. In contrast to some other studies (e.g. Teng et al., 2017; Walsh et al., 2018; Hu et al., 2019), we first identified regions showing enrichment of introgressed variants and then located the candidate regions with signatures of positive selection on the introgressed variants. This approach might have rendered our ability to identify candidates for adaptive introgression conservatively and further work is needed to identify the actual targets of selection and biological functions of the candidate genes and promoters. Nevertheless, it seems reasonable that the introgressed WL-origin variants have played a key role in the adaptation of the dominantly EL-origin sticklebacks–with a recent freshwater history in the northern Fennoscandia (Feng et al., 2022)–to the warmest and most saline part of the southern Baltic Sea.

## Conclusions

To conclude, our study of genetic introgression between two divergent stickleback lineages in the Baltic Sea demonstrates that the stabilization of hybrid genomes after admixture is a multi-stage process where the purifying selection against introgressed deleterious variations has played a central role. The varying genomic landscapes of foreign ancestry are likely the consequence of different types and targets of selection and their interactions, as well as the distribution of functional elements and the variation in recombination. Our work adds a well-worked example to studies showing that introgression can contribute to local adaptation, in spite of the widespread evidence suggesting that selection against introgression is pervasive. Although the observed weak correlation between levels of introgression and recombination rate is in stark contrast to findings in most earlier studies (e.g. Schumer et al., 2018; Edelman et al., 2019; Martin et al., 2019; Stankowski et al., 2019), it highlights the complexity of selection on shaping the genomic landscape of introgression since the occurrence of the admixture event. While more work is needed to distinguish the different forces shaping the ancestry of hybrid genomes, our findings bring new insights into the formation of a heterogeneous landscape of introgression and highlight the importance of considering demographic history, genome structure, parental population differentiation as well as recombination rate in understanding introgression.

## Author Contributions

AL and JM conceived the original idea, with significant later contributions from XF. XF and AL analyzed the data. XF took the lead in writing the manuscript, with significant contributions from AL and JM.

## Acknowledgments

We thank our collaborators and colleagues for their help in obtaining the samples (listed in Acknowledgements of Feng et al., 2022). The advice and support from Paolo Momigliano, Pasi Rastas, Petri Kemppainen, Mikko Kivikoski, Simon Martin, and Martin Petr is gratefully acknowledged. Our research was supported by grants from the Academy Finland (# 129662, 134728 and 218343 to JM; # 322681 to AL), Helsinki Lifesciences Center (HiLife; to JM), Chinese Scholarship Council (# 201608520032 to XF) and Finnish Cultural Foundation (#00210295 to XF). Computational resources provided by the CSC–IT Center for Science, Finland, are acknowledged with gratitude.

## Data availability statement

The whole-genome re-sequencing data have been published previously in Feng et al. (2022) and Feng et al. (2023). All the raw sequence data relevant to this study can be found in European Nucleotide Archive (ENA) (https://www.ebi.ac.uk/ena) through accession code PRJEB39599. Other relevant data can be found in the Zenodo Open Repository: https://zenodo.org/record/xxxxx.

## Supplementary Materials

### Supplementary Figures

Supplementary Figure S1. Four candidate regions of adaptive introgression identified with *fd*, *U20* and *Q95* analyses (Methods). The panels show the gene annotations (coding sequences in orange), per site variant allele frequencies (VAF; heatmap) for the three sets of population (WL source, BS7, EL source), and *F_ST_*, *d_xy_* and *π* (10kb-windows) in the northern Baltic Sea populations (BS7). The red and blue colors indicate *F_ST_* and *d_xy_* measured against DEN-NOR (WL source) and RUS-LEV (EL source), respectively.

Supplementary Figure S2. The *r_xy_* statistics for coding variants from five different recombination rate bins. (a) Show the relationship between recombination rate bins and *r_xy_* across pairwise comparisons (Spearman’s *rs*=−0.176, *p*=0.220), and (b) show the variation across recombination rate bins within each pairwise comparison.

Supplementary Tables

## References

1. Aldenhoven, J. T., Miller, M. A., Corneli, P. S., & Shapiro, M. D. (2010). Phylogeography of ninespine sticklebacks (*Pungitius pungitius*) in North America: 、 glacial refugia and the origins of adaptive traits. Molecular Ecology, 19(18), 4061–4076. 10.1111/j.1365-294X.2010.04801.x

2. Amorim, C. E. G., Hofer, T., Ray, N., Foll, M., Ruiz-Linares, A., & Excoffier, L. (2017). Long-distance dispersal suppresses introgression of local alleles during range expansions. Heredity, 118(2), 135–142. 10.1038/hdy.2016.68

3. Arnegard, M. E., McGee, M. D., Matthews, B., Marchinko, K. B., Conte, G. L., Kabir, S., Bedford, N., Bergek, S., Chan, Y. F., Jones, F. C., Kingsley, D. M., Peichel, C. L., & Schluter, D. (2014). Genetics of ecological divergence during speciation. Nature, 511(7509), 307–311. 10.1038/nature13301

4. Baker, Z., Schumer, M., Haba, Y., Bashkirova, L., Holland, C., Rosenthal, G. G., & Przeworski, M. (2017). Repeated losses of PRDM9-directed recombination despite the conservation of PRDM9 across vertebrates. ELife, 6, e24133. 10.7554/eLife.24133

5. Bay, R. A., Taylor, E. B., & Schluter, D. (2019). Parallel introgression and selection on introduced alleles in a native species. Molecular Ecology, 28(11), 2802–2813. 10.1111/mec.15097

6. Benjamini, Y., & Hochberg, Y. (1995). Controlling the false discovery rate: a practical and powerful approach to multiple testing. Journal of the Royal Statistical Society: Series B (Methodological), 57(1), 289–300. 10.1111/j.2517-6161.1995.tb02031.x

7. Björck, S. (2008). The late quaternary development of the Baltic Sea basin. In Assessment of climate change for the Baltic Sea Basin (pp. 398–407). Springer.

8. Bomblies, K., Lempe, J., Epple, P., Warthmann, N., Lanz, C., Dangl, J. L., & Weigel, D. (2007). Autoimmune response as a mechanism for a Dobzhansky-Muller-Type incompatibility syndrome in plants. PLoS Biology, 5(9), e236. 10.1371/journal.pbio.0050236

9. Bruneaux, M., Johnston, S. E., Herczeg, G., Merilä, J., Primmer, C. R., & Vasemägi, A. (2013). Molecular evolutionary and population genomic analysis of the nine-spined stickleback using a modified restriction-site-associated DNA tag approach. Molecular Ecology, 22(3), 565–582. 10.1111/j.1365-294X.2012.05749.x

10. Calfee, E., Gates, D., Lorant, A., Perkins, M. T., Coop, G., & Ross-Ibarra, J. (2021). Selective sorting of ancestral introgression in maize and teosinte along an elevational cline. bioRxiv. 10.1101/2021.03.05.434040

11. Christie, K., & Strauss, S. Y. (2018). Along the speciation continuum: Quantifying intrinsic and extrinsic isolating barriers across five million years of evolutionary divergence in California jewelflowers. Evolution, 72(5), 1063–1079. 10.1111/evo.13477

12. Chunco, A. J. (2014). Hybridization in a warmer world. Ecology and Evolution, 4(10), 2019–2031. 10.1002/ece3.1052

13. Cingolani, P., Platts, A., Wang, L. L., Coon, M., Nguyen, T., Wang, L., Land, S. J., Lu, X., & Ruden, D. M. (2012). A program for annotating and predicting the effects of single nucleotide polymorphisms, *SnpEff*: SNPs in the genome of Drosophila melanogaster strain w ^1118^ ; iso-2; iso-3. Fly, 6(2), 80–92. 10.4161/fly.19695

14. Corbett-Detig, R., & Nielsen, R. (2017). A hidden markov model approach for simultaneously estimating local ancestry and admixture time using next generation sequence data in samples of arbitrary ploidy. PLOS Genetics, 13(1), e1006529. 10.1371/journal.pgen.1006529

15. Danecek, P., Auton, A., Abecasis, G., Albers, C. A., Banks, E., DePristo, M. A., Handsaker, R. E., Lunter, G., Marth, G. T., Sherry, S. T., McVean, G., Durbin, R., & 1000 Genomes Project Analysis Group. (2011). The variant call format and VCFtools. Bioinformatics, 27(15), 2156–2158. 10.1093/bioinformatics/btr330

16. Dannemann, M., & Kelso, J. (2017). The contribution of Neanderthals to phenotypic variation in modern humans. The American Journal of Human Genetics, 101(4), 578–589. 10.1016/j.ajhg.2017.09.010

17. Dannemann, M., Prüfer, K., & Kelso, J. (2017). Functional implications of Neandertal introgression in modern humans. Genome Biology, 18(1), 61. 10.1186/s13059-017-1181-7

18. Dowling, T. E., & Secor, C. L. (1997). The role of hybridization and introgression in the diversification of animals. Annual Review of Ecology and Systematics, 28(1), 593–619. 10.1146/annurev.ecolsys.28.1.593

19. Duranton, M., & Pool, J. E. (2022). Interactions between natural selection and recombination shape the genomic landscape of introgression. Molecular Biology and Evolution, 39(7), msac122. 10.1093/molbev/msac122

20. Edelman, N. B., Frandsen, P. B., Miyagi, M., Clavijo, B., Davey, J., Dikow, R. B., García-Accinelli, G., Van Belleghem, S. M., Patterson, N., Neafsey, D. E., Challis, R., Kumar, S., Moreira, G. R. P., Salazar, C., Chouteau, M., Counterman, B. A., Papa, R., Blaxter, M., Reed, R. D., … Mallet, J. (2019). Genomic architecture and introgression shape a butterfly radiation. Science, 366(6465), 594–599. 10.1126/science.aaw2090

21. Edelman, N. B., & Mallet, J. (2021). Prevalence and adaptive impact of introgression. Annual Review of Genetics, 55(1), 265–283. 10.1146/annurev-genet-021821-020805

22. Feng, X., Löytynoja, A., & Merilä, J. (2023). Estimating recent and historical effective population size of marine and freshwater sticklebacks. bioRxiv. 10.1101/2023.05.22.541730

23. Feng, X., Merilä, J., & Löytynoja, A. (2022). Complex population history affects admixture analyses in nine-spined sticklebacks. Molecular Ecology, 31(20), 5386–5401. 10.1111/mec.16651

24. Frith, M. C., Hamada, M., & Horton, P. (2010). Parameters for accurate genome alignment. BMC Bioinformatics, 11(1), 80. 10.1186/1471-2105-11-80

25. Garcia, H., Weathers, K., Paver, C., Smolyar, I., Boyer, T., Locarnini, M., Zweng, M., Mishonov, A., Baranova, O., Seidov, D., & others. (2019). World Ocean Atlas 2018, Volume 3: Dissolved oxygen, apparent oxygen utilization, and dissolved oxygen saturation.

26. Gittelman, R. M., Schraiber, J. G., Vernot, B., Mikacenic, C., Wurfel, M. M., & Akey, J. M. (2016). Archaic hominin admixture facilitated adaptation to Out-of-Africa environments. Current Biology, 26(24), 3375–3382. 10.1016/j.cub.2016.10.041

27. Groh, J., & Coop, G. (2023). The temporal and genomic scale of selection following hybridization. bioRxiv. 10.1101/2023.05.25.542345

28. Guo, B., Fang, B., Shikano, T., Momigliano, P., Wang, C., Kravchenko, A., & Merilä, J. (2019). A phylogenomic perspective on diversity, hybridization and evolutionary affinities in the stickleback genus *Pungitius*. Molecular Ecology, 28(17), 4046–4064. 10.1111/mec.15204

29. Harris, K., & Nielsen, R. (2016). The Genetic cost of Neanderthal introgression. Genetics, 203(2), 881–891. 10.1534/genetics.116.186890

30. Harrison, R. G., & Larson, E. L. (2014). Hybridization, introgression, and the nature of species boundaries. Journal of Heredity, 105(S1), 795–809. 10.1093/jhered/esu033

31. Hedrick, P. W. (2013). Adaptive introgression in animals: Examples and comparison to new mutation and standing variation as sources of adaptive variation. Molecular Ecology, 22(18), 4606–4618. 10.1111/mec.12415

32. Herczeg, G., Gonda, A., & Merilä, J. (2009). Evolution of gigantism in nine-spined sticklebacks. Evolution, 63(12), 3190–3200. 10.1111/j.1558-5646.2009.00781.x

33. Herczeg, G., Turtiainen, M., & Merilä, J. (2010). Morphological divergence of North-European nine-spined sticklebacks (*Pungitius pungitius*): Signatures of parallel evolution. Biological Journal of the Linnean Society, 101(2), 403–416. 10.1111/j.1095-8312.2010.01518.x

34. Herrero, J., Muffato, M., Beal, K., Fitzgerald, S., Gordon, L., Pignatelli, M., Vilella, A. J., Searle, S. M. J., Amode, R., Brent, S., Spooner, W., Kulesha, E., Yates, A., & Flicek, P. (2016). Ensembl comparative genomics resources. Database, 2016, bav096. 10.1093/database/bav096

35. Hu, X.-J., Yang, J., Xie, X.-L., Lv, F.-H., Cao, Y.-H., Li, W.-R., Liu, M.-J., Wang, Y.-T., Li, J.-Q., Liu, Y.-G., Ren, Y.-L., Shen, Z.-Q., Wang, F., Hehua, Ee., Han, J.-L., & Li, M.-H. (2019). The genome landscape of Tibetan sheep reveals adaptive introgression from argali and the history of early human settlements on the Qinghai–Tibetan plateau. Molecular Biology and Evolution, 36(2), 283–303. 10.1093/molbev/msy208

36. Huerta-Sánchez, E., Jin, X., Asan, Bianba, Z., Peter, B. M., Vinckenbosch, N., Liang, Y., Yi, X., He, M., Somel, M., Ni, P., Wang, B., Ou, X., Huasang, Luosang, J., Cuo, Z. X. P., Li, K., Gao, G., Yin, Y., … Nielsen, R. (2014). Altitude adaptation in Tibetans caused by introgression of Denisovan-like DNA. Nature, 512(7513), 194–197. 10.1038/nature13408

37. Jagoda, E., Lawson, D. J., Wall, J. D., Lambert, D., Muller, C., Westaway, M., Leavesley, M., Capellini, T. D., Mirazón Lahr, M., Gerbault, P., Thomas, M. G., Migliano, A. B., Willerslev, E., Metspalu, M., & Pagani, L. (2018). Disentangling immediate adaptive introgression from selection on standing introgressed variation in humans. Molecular Biology and Evolution, 35(3), 623–630. 10.1093/molbev/msx314

38. Jokinen, H., Momigliano, P., & Merilä, J. (2019). From ecology to genetics and back: The tale of two flounder species in the Baltic Sea. ICES Journal of Marine Science, 76(7), 2267–2275. 10.1093/icesjms/fsz151

39. Jovine, L., Darie, C. C., Litscher, E. S., & Wassarman, P. M. (2005). Zona pellucida domain proteins. Annual Review of Biochemistry, 74(1), 83–114. 10.1146/annurev.biochem.74.082803.133039

40. Juric, I., Aeschbacher, S., & Coop, G. (2016). The strength of selection against Neanderthal introgression. PLOS Genetics, 12(11), e1006340. 10.1371/journal.pgen.1006340

41. Kim, B. Y., Huber, C. D., & Lohmueller, K. E. (2018). Deleterious variation shapes the genomic landscape of introgression. PLOS Genetics, 14(10), e1007741. 10.1371/journal.pgen.1007741

42. Kivikoski, M., Rastas, P., Löytynoja, A., & Merilä, J. (2021). Automated improvement of stickleback reference genome assemblies with Lep-Anchor software. Molecular Ecology Resources, 21(6), 2166–2176. 10.1111/1755-0998.13404

43. Kovach, R. P., Hand, B. K., Hohenlohe, P. A., Cosart, T. F., Boyer, M. C., Neville, H. H., Muhlfeld, C. C., Amish, S. J., Carim, K., Narum, S. R., Lowe, W. H., Allendorf, F. W., & Luikart, G. (2016). Vive la résistance: Genome-wide selection against introduced alleles in invasive hybrid zones. Proceedings of the Royal Society B: Biological Sciences, 283(1843), 20161380. 10.1098/rspb.2016.1380

44. Kuhlwilm, M., Gronau, I., Hubisz, M. J., De Filippo, C., Prado-Martinez, J., Kircher, M., Fu, Q., Burbano, H. A., Lalueza-Fox, C., De La Rasilla, M., Rosas, A., Rudan, P., Brajkovic, D., Kucan, Ž., Gušic, I., Marques-Bonet, T., Andrés, A. M., Viola, B., Pääbo, S., … Castellano, S. (2016). Ancient gene flow from early modern humans into Eastern Neanderthals. Nature, 530(7591), 429–433. 10.1038/nature16544

45. Kyriazis, C. C., Wayne, R. K., & Lohmueller, K. E. (2021). Strongly deleterious mutations are a primary determinant of extinction risk due to inbreeding depression. Evolution Letters, 5(1), 33–47. 10.1002/evl3.209

46. Lamichhaney, S., Han, F., Webster, M. T., Andersson, L., Grant, B. R., & Grant, P. R. (2018). Rapid hybrid speciation in Darwin’s finches. Science, 359(6372), 224–228. 10.1126/science.aao4593

47. Lee, H.-Y., Chou, J.-Y., Cheong, L., Chang, N.-H., Yang, S.-Y., & Leu, J.-Y. (2008). Incompatibility of nuclear and mitochondrial genomes causes hybrid sterility between two yeast species. Cell, 135(6), 1065–1073. 10.1016/j.cell.2008.10.047

48. Li, H. (2013). Aligning sequence reads, clone sequences and assembly contigs with BWA-MEM. ArXiv Preprint ArXiv:1303.3997.

49. Li, H., Handsaker, B., Wysoker, A., Fennell, T., Ruan, J., Homer, N., Marth, G., Abecasis, G., Durbin, R., & 1000 Genome Project Data Processing Subgroup. (2009). The sequence alignment/map format and SAMtools. Bioinformatics, 25(16), 2078–2079. 10.1093/bioinformatics/btp352

50. Liu, S., Zhang, L., Sang, Y., Lai, Q., Zhang, X., Jia, C., Long, Z., Wu, J., Ma, T., Mao, K., Street, N. R., Ingvarsson, P. K., Liu, J., & Wang, J. (2022). Demographic history and natural selection shape patterns of deleterious mutation load and barriers to introgression across *Populus* genome. Molecular Biology and Evolution, 39(2), msac008. 10.1093/molbev/msac008

51. Lønning, S., & Solemdal, P. (1972). The relation between thickness of chorion and specific gravity of eggs from Norwegian and Baltic flatfish populations.

52. Malinsky, M., Svardal, H., Tyers, A. M., Miska, E. A., Genner, M. J., Turner, G. F., & Durbin, R. (2018). Whole-genome sequences of Malawi cichlids reveal multiple radiations interconnected by gene flow. Nature Ecology & Evolution, 2(12), 1940–1955. 10.1038/s41559-018-0717-x

53. Mallet, J. (2005). Hybridization as an invasion of the genome. Trends in Ecology & Evolution, 20(5), 229–237. 10.1016/j.tree.2005.02.010

54. Marques, D. A., Lucek, K., Sousa, V. C., Excoffier, L., & Seehausen, O. (2019). Admixture between old lineages facilitated contemporary ecological speciation in Lake Constance stickleback. Nature Communications, 10(1), 4240. 10.1038/s41467-019-12182-w

55. Martin, S. H., Davey, J. W., & Jiggins, C. D. (2015). Evaluating the use of ABBA–BABA statistics to locate introgressed loci. Molecular Biology and Evolution, 32(1), 244–257. 10.1093/molbev/msu269

56. Martin, S. H., Davey, J. W., Salazar, C., & Jiggins, C. D. (2019). Recombination rate variation shapes barriers to introgression across butterfly genomes. PLOS Biology, 17(2), e2006288. 10.1371/journal.pbio.2006288

57. Martin, S. H., & Jiggins, C. D. (2017). Interpreting the genomic landscape of introgression. Current Opinion in Genetics & Development, 47, 69–74. 10.1016/j.gde.2017.08.007

58. Masly, J. P., & Presgraves, D. C. (2007). High-resolution genome-wide dissection of the two rules of speciation in *Drosophila*. PLoS Biology, 5(9), e243. 10.1371/journal.pbio.0050243

59. Matute, D. R., Comeault, A. A., Earley, E., Serrato-Capuchina, A., Peede, D., Monroy-Eklund, A., Huang, W., Jones, C. D., Mackay, T. F. C., & Coyne, J. A. (2020). Rapid and predictable evolution of admixed populations between two *Drosophila* species pairs. Genetics, 214(1), 211–230. 10.1534/genetics.119.302685

60. McCoy, R. C., Wakefield, J., & Akey, J. M. (2017). Impacts of Neanderthal-introgressed sequences on the landscape of human gene expression. Cell, 168(5), 916–927.e12. 10.1016/j.cell.2017.01.038

61. McKenna, A., Hanna, M., Banks, E., Sivachenko, A., Cibulskis, K., Kernytsky, A., Garimella, K., Altshuler, D., Gabriel, S., Daly, M., & DePristo, M. A. (2010). The Genome Analysis Toolkit: A MapReduce framework for analyzing next-generation DNA sequencing data. Genome Research, 20(9), 1297–1303. 10.1101/gr.107524.110

62. Meier, J. I., Marques, D. A., Mwaiko, S., Wagner, C. E., Excoffier, L., & Seehausen, O. (2017). Ancient hybridization fuels rapid cichlid fish adaptive radiations. Nature Communications, 8(1), 14363. 10.1038/ncomms14363

63. Momigliano, P., Jokinen, H., Fraimout, A., Florin, A.-B., Norkko, A., & Merilä, J. (2017). Extraordinarily rapid speciation in a marine fish. Proceedings of the National Academy of Sciences, 114(23), 6074–6079. 10.1073/pnas.1615109114

64. Moran, B. M., Payne, C., Langdon, Q., Powell, D. L., Brandvain, Y., & Schumer, M. (2021). The genomic consequences of hybridization. ELife, 10, e69016. 10.7554/eLife.69016

65. Myers, S., Bowden, R., Tumian, A., Bontrop, R. E., Freeman, C., MacFie, T. S., McVean, G., & Donnelly, P. (2010). Drive against hotspot motifs in primates implicates the *PRDM9* gene in meiotic recombination. Science, 327(5967), 876–879. 10.1126/science.1182363

66. Nachman, M. W., & Payseur, B. A. (2012). Recombination rate variation and speciation: Theoretical predictions and empirical results from rabbits and mice. Philosophical Transactions of the Royal Society B: Biological Sciences, 367(1587), 409–421. 10.1098/rstb.2011.0249

67. Natri, H. M., Merilä, J., & Shikano, T. (2019). The evolution of sex determination associated with a chromosomal inversion. Nature Communications, 10(1), 145. 10.1038/s41467-018-08014-y

68. Nissling, A. (2002). Reproductive success in relation to salinity for three flatfish species, dab (*Limanda limanda*), plaice (*Pleuronectes platessa*), and flounder (*Pleuronectes flesus*), in the brackish water Baltic Sea. ICES Journal of Marine Science, 59(1), 93–108. 10.1006/jmsc.2001.1134

69. Oziolor, E. M., Reid, N. M., Yair, S., Lee, K. M., Guberman VerPloeg, S., Bruns, P. C., Shaw, J. R., Whitehead, A., & Matson, C. W. (2019). Adaptive introgression enables evolutionary rescue from extreme environmental pollution. Science, 364(6439), 455–457. 10.1126/science.aav4155

70. Palumbi, S. R. (1994). Genetic divergence, reproductive isolation, and marine speciation. Annual Review of Ecology and Systematics, 25(1), 547–572. 10.1146/annurev.es.25.110194.002555

71. Pardo-Diaz, C., Salazar, C., Baxter, S. W., Merot, C., Figueiredo-Ready, W., Joron, M., McMillan, W. O., & Jiggins, C. D. (2012). Adaptive introgression across species boundaries in *Heliconius* butterflies. PLoS Genetics, 8(6), e1002752. 10.1371/journal.pgen.1002752

72. Patterson, N., Moorjani, P., Luo, Y., Mallick, S., Rohland, N., Zhan, Y., Genschoreck, T., Webster, T., & Reich, D. (2012). Ancient admixture in human history. Genetics, 192(3), 1065–1093. 10.1534/genetics.112.145037

73. Petr, M., Pääbo, S., Kelso, J., & Vernot, B. (2019). Limits of long-term selection against Neandertal introgression. Proceedings of the National Academy of Sciences, 116(5), 1639–1644. 10.1073/pnas.1814338116

74. Pool, J. E. (2015). The mosaic ancestry of the *Drosophila* genetic reference panel and the *D. melanogaster* reference genome reveals a network of epistatic fitness interactions. Molecular Biology and Evolution, msv194. 10.1093/molbev/msv194

75. Racimo, F., Marnetto, D., & Huerta-Sánchez, E. (2016). Signatures of archaic adaptive introgression in present-day human populations. Molecular Biology and Evolution, msw216. 10.1093/molbev/msw216

76. Racimo, F., Sankararaman, S., Nielsen, R., & Huerta-Sánchez, E. (2015). Evidence for archaic adaptive introgression in humans. Nature Reviews Genetics, 16(6), 359–371. 10.1038/nrg3936

77. Reich, D., Thangaraj, K., Patterson, N., Price, A. L., & Singh, L. (2009). Reconstructing Indian population history. Nature, 461(7263), 489–494. 10.1038/nature08365

78. Reusch, T. B. H., Dierking, J., Andersson, H. C., Bonsdorff, E., Carstensen, J., Casini, M., Czajkowski, M., Hasler, B., Hinsby, K., Hyytiäinen, K., Johannesson, K., Jomaa, S., Jormalainen, V., Kuosa, H., Kurland, S., Laikre, L., MacKenzie, B. R., Margonski, P., Melzner, F., … Zandersen, M. (2018). The Baltic Sea as a time machine for the future coastal ocean. Science Advances, 4(5), eaar8195. 10.1126/sciadv.aar8195

79. Sankararaman, S., Mallick, S., Dannemann, M., Prüfer, K., Kelso, J., Pääbo, S., Patterson, N., & Reich, D. (2014). The genomic landscape of Neanderthal ancestry in present-day humans. Nature, 507(7492), 354–357. 10.1038/nature12961

80. Sankararaman, S., Mallick, S., Patterson, N., & Reich, D. (2016). The combined landscape of Denisovan and Neanderthal ancestry in present-day humans. Current Biology, 26(9), 1241–1247. 10.1016/j.cub.2016.03.037

81. Schumer, M., Xu, C., Powell, D. L., Durvasula, A., Skov, L., Holland, C., Blazier, J. C., Sankararaman, S., Andolfatto, P., Rosenthal, G. G., & Przeworski, M. (2018). Natural selection interacts with recombination to shape the evolution of hybrid genomes. Science, 360(6389), 656–660. 10.1126/science.aar3684

82. Setter, D., Mousset, S., Cheng, X., Nielsen, R., DeGiorgio, M., & Hermisson, J. (2020). VolcanoFinder: Genomic scans for adaptive introgression. PLOS Genetics, 16(6), e1008867. 10.1371/journal.pgen.1008867

83. Shikano, T., Ramadevi, J., & Merila, J. (2010). Identification of local- and habitat-dependent selection: Scanning functionally important genes in nine-spined sticklebacks (*Pungitius pungitius*). Molecular Biology and Evolution, 27(12), 2775–2789. 10.1093/molbev/msq167

84. Shikano, T., Shimada, Y., Herczeg, G., & Merilä, J. (2010). History vs. habitat type: Explaining the genetic structure of European nine-spined stickleback (*Pungitius pungitius*) populations. Molecular Ecology, 19(6), 1147–1161. 10.1111/j.1365-294X.2010.04553.x

85. Shumate, A., & Salzberg, S. L. (2021). Liftoff: Accurate mapping of gene annotations. Bioinformatics, 37(12), 1639–1643. 10.1093/bioinformatics/btaa1016

86. Skoglund, P., Ersmark, E., Palkopoulou, E., & Dalén, L. (2015). Ancient wolf genome reveals an early divergence of domestic dog ancestors and admixture into high-latitude breeds. Current Biology, 25(11), 1515–1519. 10.1016/j.cub.2015.04.019

87. Smukowski Heil, C. S., Large, C. R. L., Patterson, K., Hickey, A. S.-M., Yeh, C.-L. C., & Dunham, M. J. (2019). Temperature preference can bias parental genome retention during hybrid evolution. PLOS Genetics, 15(9), e1008383. 10.1371/journal.pgen.1008383

88. Stankowski, S., Chase, M. A., Fuiten, A. M., Rodrigues, M. F., Ralph, P. L., & Streisfeld, M. A. (2019). Widespread selection and gene flow shape the genomic landscape during a radiation of monkeyflowers. PLOS Biology, 17(7), e3000391. 10.1371/journal.pbio.3000391

89. Suarez-Gonzalez, A., Lexer, C., & Cronk, Q. C. B. (2018). Adaptive introgression: A plant perspective. Biology Letters, 14(3), 20170688. 10.1098/rsbl.2017.0688

90. Teacher, A. G. F., Shikano, T., Karjalainen, M. E., & Merilä, J. (2011). Phylogeography and genetic structuring of European nine-spined sticklebacks (*Pungitius pungitius*)—mitochondrial DNA evidence. PLoS ONE, 6(5), e19476. 10.1371/journal.pone.0019476

91. Telis, N., Aguilar, R., & Harris, K. (2020). Selection against archaic hominin genetic variation in regulatory regions. Nature Ecology & Evolution, 4(11), 1558–1566. 10.1038/s41559-020-01284-0

92. Teng, H., Zhang, Y., Shi, C., Mao, F., Cai, W., Lu, L., Zhao, F., Sun, Z., & Zhang, J. (2017). Population genomics reveals speciation and introgression between brown Norway rats and their sibling species. Molecular Biology and Evolution, 34(9), 2214–2228. 10.1093/molbev/msx157

93. The Heliconius Genome Consortium. (2012). Butterfly genome reveals promiscuous exchange of mimicry adaptations among species. Nature, 487(7405), 94–98. 10.1038/nature11041

94. Tigano, A., & Friesen, V. L. (2016). Genomics of local adaptation with gene flow. Molecular Ecology, 25(10), 2144–2164. 10.1111/mec.13606

95. Turner, R. M. (2003). Tales From the Tail: What do we really know about sperm motility? Journal of Andrology, 24(6), 790–803. 10.1002/j.1939-4640.2003.tb03123.x

96. Tyner, C., Barber, G. P., Casper, J., Clawson, H., Diekhans, M., Eisenhart, C., Fischer, C. M., Gibson, D., Gonzalez, J. N., Guruvadoo, L., Haeussler, M., Heitner, S., Hinrichs, A. S., Karolchik, D., Lee, B. T., Lee, C. M., Nejad, P., Raney, B. J., Rosenbloom, K. R., … Kent, W. J. (2017). The UCSC Genome Browser database: 2017 update. Nucleic Acids Research, 45(Database issue), D626–D634. 10.1093/nar/gkw1134

97. Ukkonen, P., Aaris-Sørensen, K., Arppe, L., Daugnora, L., Halkka, A., Lõugas, L., Oinonen, M. J., Pilot, M., & Storå, J. (2014). An Arctic seal in temperate waters: History of the ringed seal (*Pusa hispida*) in the Baltic Sea and its adaptation to the changing environment. The Holocene, 24(12), 1694–1706. 10.1177/0959683614551226

98. Varadharajan, S., Rastas, P., Löytynoja, A., Matschiner, M., Calboli, F. C. F., Guo, B., Nederbragt, A. J., Jakobsen, K. S., & Merilä, J. (2019). A high-quality assembly of the nine-spined stickleback (*Pungitius pungitius*) genome. Genome Biology and Evolution, 11(11), 3291–3308. 10.1093/gbe/evz240

99. Vattathil, S., & Akey, J. M. (2015). Small amounts of archaic admixture provide big insights into human history. Cell, 163(2), 281–284. 10.1016/j.cell.2015.09.042

100. Veller, C., Edelman, N. B., Muralidhar, P., & Nowak, M. A. (2023). Recombination and selection against introgressed DNA. Evolution, 77(4), 1131–1144. 10.1093/evolut/qpad021

101. Walsh, J., Kovach, A. I., Olsen, B. J., Shriver, W. G., & Lovette, I. J. (2018). Bidirectional adaptive introgression between two ecologically divergent sparrow species. Evolution, 72(10), 2076–2089. 10.1111/evo.13581

102. Wang, Y., Wang, Y., Cheng, X., Ding, Y., Wang, C., Merilä, J., & Guo, B. (2023). Prevalent introgression underlies convergent evolution in the diversification of *Pungitius* sticklebacks. Molecular Biology and Evolution, 40(2), msad026. 10.1093/molbev/msad026

103. Whitney, K. D., Randell, R. A., & Rieseberg, L. H. (2006). Adaptive introgression of herbivore resistance traits in the weedy sunflower *Helianthus annuus*. The American Naturalist, 167(6), 794–807. 10.1086/504606

104. Wu, C.-I. (2001). The genic view of the process of speciation: Genic view of the process of speciation. Journal of Evolutionary Biology, 14(6), 851–865. 10.1046/j.1420-9101.2001.00335.x

105. Xue, Y., Prado-Martinez, J., Sudmant, P. H., Narasimhan, V., Ayub, Q., Szpak, M., Frandsen, P., Chen, Y., Yngvadottir, B., Cooper, D. N., De Manuel, M., Hernandez-Rodriguez, J., Lobon, I., Siegismund, H. R., Pagani, L., Quail, M. A., Hvilsom, C., Mudakikwa, A., Eichler, E. E., … Scally, A. (2015). Mountain gorilla genomes reveal the impact of long-term population decline and inbreeding. Science, 348(6231), 242–245. 10.1126/science.aaa3952

106. Zhang, W., Dasmahapatra, K. K., Mallet, J., Moreira, G. R. P., & Kronforst, M. R. (2016). Genome-wide introgression among distantly related Heliconius butterfly species. Genome Biology, 17(1), 25. 10.1186/s13059-016-0889-0

107. Zuccarello, D., Ferlin, A., Cazzadore, C., Pepe, A., Garolla, A., Moretti, A., Cordeschi, G., Francavilla, S., & Foresta, C. (2008). Mutations in dynein genes in patients affected by isolated non-syndromic asthenozoospermia. Human Reproduction, 23(8), 1957–1962. 10.1093/humrep/den193

